# Categorization of prophage genes in *Bacillus subtilis* 168 and assessing their relative importance through RNA-seq gene expression analysis

**DOI:** 10.1101/2021.10.26.466030

**Authors:** Wenfa Ng

## Abstract

Bacteriophage evolves to control the population of fast-growing bacterial cells, without which explosion in bacterial population may induce unimaginable harm to diverse ecosystems. But, bacteriophage also “hide” in bacterial genomes when nutritional and environmental circumstances are unfavourable. This involves the integration of phage genome into the host genome at appropriate genomic loci in a process known as lysogeny. This work sought to delineate the prophages present in the annotated genome of *Bacillus subtilis* 168, and assess their relative importance through RNA-seq expression analysis. Firstly, examination of the annotated genome of the model Gram-positive bacterium revealed five distinct prophage regions: SPBeta, prophage 6, PBSX, prophage 3 region, and prophage 1 region. All prophage regions contain host genes, which suggests that host transposase activity have swapped in host genes for phage genes in the prophage genome. Given the significant number of phage genes that have been swapped into each of the prophage genome, all prophage regions are deemed to be defective. BLAST analysis further highlighted that many of the prophages in *B. subtilis* are extinct given that they do not have ancestral or daughter brethren. However, RNA-seq transcriptome analysis of *B. subtilis* turned out an interesting paradox indicative of the important role that host transposase have in swapping in host promoters for prophage genes. Specifically, a significant number of prophage genes are highly expressed, which is implausible given that phage genes should be transcriptionally silent. The result and phenomenon further suggests the relative facile nature in which host promoters could be swapped in for phage genes, which is indicative of presence of genomic motifs in prophage genome recognizable by host transposase. Existence of such sequence motifs is thus indicative of possible co-evolution of transposase and phages where transposases were originally a part of the phage genome, which latter “jumped out” into the host genome to aid the swapping in of host genes into the prophage genome for augmenting prophage genetic repertoire in the face of changing environmental conditions. Overall, it is not uncommon for bacterial species to harbour multiple prophages. But, lysogeny may not be a viable option for long-term preservation of prophage genetic repertoire given that host transposase would inevitable swap in host genes at random locations in the prophage genome.

## 1. Introduction

Bacteriophages play a hugely important role in microbial ecology as they help check the population of bacterial species in diverse habitats. By constraining the population of bacterial species, phages prevent the excessive growth of particular bacterial species that may threaten the survival of other microbial species. This then ensures ecosystem homeostasis which is significant as it safeguards the ecological functions that microbial species bring to Earth.

Although it is known that bacteriophages infect specific bacterial species through receptor mediated binding, there remains significant challenges in identifying them and studying them in bacterial populations. One key challenge is in timing the lytic cycle of a prophage.^1^ Typically, bacteriophages enter the lysogenic mode when circumstances in the environment are unfavourable for their survival.^2 3^ This phase of the phage life cycle entails the integration of the entire phage genome into the bacterial genome.^4^ Bacteriophages that have activated lysogeny and whose genome has been integrated into the host genome are known as prophages.

Prophages can be viewed as bacteriophage in quiescent mode as they remain fully capable of entering the lytic phase where the whole phage genome is excised from the host genome, activated with phage genes transcribed, and help in the assembly of full phages able to leave and infect other bacterial cells of the same species. But, lysogeny also presents risk to the phage. Chief of which is the irresistible effect of evolutionary pressure on the integrity of the phage genes. In this scenario, phage genes in the host genome would remain transcriptionally silent, and thus given that they do not encode functional protein of use to the host cell, there is no driver that could guard against random genetic drift resulting from mutations incurred during DNA replication of the host genome. Hence, phage genes integrated into host genome may suffer from genetic drift, and through a relatively short span of evolutionary time, lose their functionality. If sufficient number of phage genes suffer from this fate, the prophage may lose its ability to leave the host cell, and is thus inactivated.

But, a more pressing challenge to the survival of a prophage in lysogeny mode is the threat from transposon. These small genetic elements activated by transposase could copy a functional gene from the host into genomic loci occupied by phage genes. Such random insertion events thus disrupt phage genes, and would remain retained by the host cells, as loss of phage genes do not affect the fitness of the host cell. Hence, cells with high level of transposase and transposons do not present favourable environment for the long-term survival of prophages, with the end result that these bacterial species would usually have prophage with truncated genome or which carry genes that belongs to the host rather than the phage itself.

Traditionally, research in bacteriophage genetics have relied on Sanger sequencing for analyzing each gene in turn, which are subsequently cloned into an expression vector for introduction to a recombinant host to produce the target protein. What follows next is biochemical assays for assessing and ascribing function to the target protein. But, this mode of meticulous and tedious phage research has increasingly been replaced by modern genome sequencing efforts, that through automated annotation pipelines, help identify many conserved hypothetical phage genes in genomes of bacterial species. Such approaches significantly streamlined the phage identification exercise, as functions of many genes in the sequenced genome of the bacterial species could be automatically annotated. Finding phage genes would then involve a relatively simple keyword search for “phage” in the Excel file containing the annotated genome of the bacterial species.

This work sought to use the above described approach in understanding the prophages present in the annotated genome of *Bacillus subtilis* 168. Specifically, an in-house MATLAB software was used in parsing the annotated genome of *B. subtilis* 168, and helped in the construction of a gene database of the microorganism. Information included in the gene database include gene identifier, gene function, and gene sequence. This database was subsequently searched with the keyword “phage” to yield a list of prophage genes in the genome of *B. subtilis* 168. Manual curation was then employed to categorized different sets of phage genes as those belonging to different prophages. Further refinement was also made to identify different functional genes such as those involved in DNA replication, transcription, repair and structural roles in each set of prophage genes. Finally, RNA-seq transcriptome data for *B. subtilis* 168 under different nutritional and environmental conditions was analyzed to yield a picture of the expression level of different phage genes under different conditions.

Results revealed the presence of five prophages in *B. subtilis* 168 genome. Many of the prophages are functionally inactivated given that many of their genes have been disrupted by transposons through either swapping in of host genes or insertion of insertion sequences (IS). More importantly, some of the prophages only retained a truncated genome, which attest to the power of evolutionary forces in obliterating many of the phage genes through time. Prophages with such truncated genome are thus also likely to have been the oldest in the *B. subtilis* genome. Functional chacterization of phage genes in each prophage genome turns out an incomplete list of genes for each function. Importantly, few prophages carried the genes belonging to the full complement of functions needed for a full functional prophage. This again provides additional evidence that many of the prophages are functionally inactive and could not enter the lytic cycle. Finally, expression analysis of phage genes revealed high expression level of phage genes that have been swapped in with the host genes. In turn, these genes behave as host genes armed with their own promoters. Further analysis of phage genes with no or low expression level likely highlights the remaining true prophage genes in the genome of the host cell.

## 2. Materials and Methods

### 2.1 Reading of annotated genome file from Genbank

Annotated genome file for *Bacillus subtilis* 168 was downloaded from Genbank (NCBI Reference Sequence: NC_000964.3). An in-house MATLAB software was used to read and parse information in the annotated genome file into an Excel gene database comprising gene name, gene function, and gene sequence for each gene in the genome of the bacterium.

### 2.2 Identification and categorization of phage genes in *Bacillus subtilis*

A keyword search for “phage” was performed using the in-built Excel search function. All genes whose gene function contain the keyword “phage” would be identified, and collected as an ensemble of phage genes in *B. subtilis* in a separate database. More importantly, the obtained ensemble of phage genes in *B. subtilis* was further curated manually to yield separate list of genes for individual prophages residing in the genome of the bacterium. This meant that a list of phage genes for each prophage in *B. subtilis* 168 is available in the new database obtained by this work.

### 2.3 Functional categorization of phage genes in each prophage

To further understand the functional roles of each phage gene in the genome of individual prophage, functional categorization was performed based on the description of each phage gene to yield a dataset comprising genes belonging to different categories such as DNA replication, transcription, structural genes, and miscellaneous.

### 2.4 RNA-seq transcriptome analysis of phage genes in *B. subtilis*

RNA-seq transcriptome of *B. subtilis* under different environmental and nutritional conditions were used to assess whether growth medium or culture conditions influenced the expression level of phage genes. Briefly, RNA-seq transcriptome of *B. subtilis* was downloaded from Array Express and processed by an in-house MATLAB RNA-seq transcriptome processing software to yield expression count for each gene in the genome of *B. subtilis* under specific nutritional and environmental conditions. During transcriptome processing, 5 bases were truncated from the 5’ and 3’ end of each sequenced read. This preprocessed sequenced read was subsequently aligned to each gene in the genome of *B. subtilis*. If the alignment was successful, the expression count of the gene would be increased by 1. This process was repeated iteratively for a total of 1.2 million sequenced reads to yield a readout of the transcriptional landscape of *B. subtilis*. The Excel file containing the output comprises gene name, gene function, gene sequence, and expression count. This file was searched with the keyword “phage” in Excel to yield a set of phage genes with expression count, which was compiled as a separate database.

## 3. Results and Discussion

### 3.1 Genetic repertoire of SPBeta prophage in *Bacillus subtilis* 168

Phage SPBeta comprises 23 genes including a serine-type phage integrase (sprA). Many of the phage genes are conserved hypothetic protein with unknown functions. Of interest is that some of the phage genes are likely to have been swapped by host genes through the action of transposon. Examples of this include: N-acetyl transferase (yokL), and aminoglycoside N3’ acetyltransferase (aacD). Interestingly, phage SPBeta comes with calcium dependent DNA nuclease, RNase, and phage toxin. Overall, lack of understanding of the functions of most of the genes associated with SPBeta precludes gaining a further appreciation of the mode of operation and function of this prophage in *B. subtilis*.

Functional categorization of the annotated genes in phage SPBeta reveals only two integral genes important to phage DNA replication (uvrX) and phage integration into the host genome (sprA) (Table 2). Interestingly, the analysis also reveals the swapping of host genes into the prophage genomes. Examples of such genes are type 1 toxin-antitoxin system (bsrG), putative N-acetyltransferase (yokL), putative Rnase (yokI), putative transposase (yoyK), a calcium-dependent DNA nuclease lipoprotein (nukF), and an aminoglycoside N3’-acetyltransferase (aacD). Overall, many of the genes in prophage SPBeta have been swapped with host genes likely due to transposase activity. Given the loss of many prophage genes, it is likely that prophage SPBeta is defective, and could not enter the lytic cycle.

**Table 1:** Genetic repertoire of SPBeta prophage in *Bacillus subtilis* 168.

| Gene | Gene function |
| --- | --- |
| sunA | sublancin 168 lantibiotic antimicrobial precursor peptide |
| uvrX | lesion bypass phage DNA polymerase |
| yolD | conserved hypothetical protein |
| yolC | conserved phage protein of unknown function |
| yolB | conserved protein of unknown function |
| yolA | conserved exported protein of unknown function |
| bsrG | phage toxin; type I toxin-antitoxin system |
| yokL | putative N-acetyltransferase |
| yokK | conserved protein of unknown function |
| yokJ | conserved protein of unknown function |
| yokI | putative RNase |
| yokH | conserved protein of unknown function |
| yoyK | fragment of putative transposase |
| yokG | conserved protein of unknown function |
| nukF | calcium-dependent DNA nuclease, lipoprotein |
| yokE | conserved protein of unknown function |
| aacD | aminoglycoside N3'-acetyltransferase |
|  | hypothetical protein |
|  | hypothetical protein |
| yokC | conserved hypothetical protein |
| yokB | hypothetical protein |
| sprA | serine-type phage integrase |

**Table 2:** Categorization of function of annotated genes in phage SPBeta in *Bacillus subtilis*.

| **Function** | **DNA replication** |
| --- | --- |
| Gene | Gene function |
| uvrX | lesion bypass phage DNA polymerase |
| sprA | serine-type phage integrase |

| **Function** | **Miscellaneous** |
| --- | --- |
| Gene | Gene function |
| bsrG | phage toxin; type I toxin-antitoxin system |
| yokL | putative N-acetyltransferase |
| yokI | putative Rnase |
| yoyK | fragment of putative transposase |
| nukF | calcium-dependent DNA nuclease, lipoprotein |
| aacD | aminoglycoside N3'-acetyltransferase |

### 3.2 Genetic repertoire of prophage 6 in the genome of *Bacillus subtilis*

Similar to prophage SPBeta, prophage 6 in *Bacillus subtilis* comprises a collection of 21 genes (Table 3). Although many of the genes are putative prophage genes or prophage related, there exists instances in which prophage genes have been swapped by host genes through the action of transposase. Example of such swapped-in host genes is the putative SOS response associated protein (yoaM). But such examples are rare, and the genome of prophage 6 consist of mainly phage related genes. This suggests that the genome of prophage 6 lacks genomic sites that could be recognized by transposase for the insertion of transposons, and thus, the genome of prophage 6 remains largely intact.

**Table 3:** Genetic repertoire of prophage 6 in *Bacillus subtilis* 168.

| **Gene** | **Gene function** |
| --- | --- |
| yoaM | putative SOS response associated protein |
| yozS | putative permease, phage-related |
| yozT | conserved hypothetical protein (putative phage origin) |
| yobB | putative transcriptional regulator from prophage |
| yozV | putative phage protein |
| yobD | transcriptional regulator (phage-related, Xre family) |
| yozH | hypothetical protein; putative defective prophage 6 |
| yozI | conserved hypothetical protein |
| yobE | putative SOS response associated phage protein |
| yozW | hypothetical protein |
| yozX | putative phage protein |
| yozY | putative transcriptional regulator from bacteriophage |
| yobHc | fragment of putative DNA phage repair protein |
| yobHm | putative phage DNA repair protein fragment |
| yozL | conserved hypothetical protein of phage origin |
| yozM | putative bacteriophage protein |
| yobI | putative phage repair NTPase with transmembrane helix |
| yoyA | putative fragment of phage protein |
| yobJ | conserved protein of unknown function |
| rttL | phage toxin ribonuclease |
| yobM | putative phage protein |
| yobO | putative phage-related pre-neck appendage |

Functional categorization of prophage genes in prophage 6 of *B. subtilis* reveals that the prophage is endowed with important genes and functions in DNA replication, transcription, and other miscellaneous functions (Table 4). For example, in the area of DNA replication, prophage 6 has three DNA repair protein genes, which suggest that the genome of prophage 6 may be vulnerable to increased rate of mutations due to an error-prone DNA replication machinery. At the level of transcription, prophage 6 is also endowed with three transcriptional regulator genes which indicates that the prophage plays an active role in regulating the transcription of its genes, thereby, providing it with a fitness advantage in timing its replication and exit from the cell after the lysogeny period. Besides DNA replication and transcription, prophage 6 is also endowed with two genes associated with DNA damage SOS response, which provides tantalizing evidence that bacteriophage also uses a mechanism similar to SOS to maintain the integrity of its genome. Finally, other phage genes with important functions include the phage related permease (yozS), phage toxin ribonuclease (rttL), and phage-related pre-neck appendage (yobO).

**Table 4:**
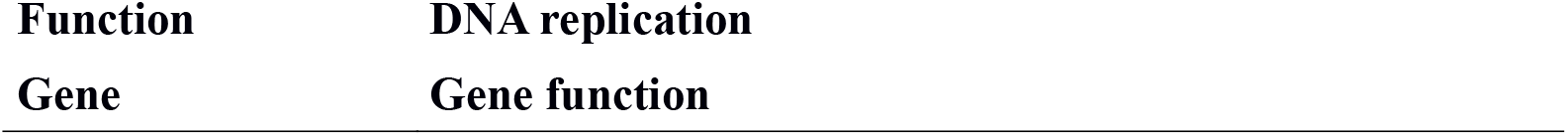
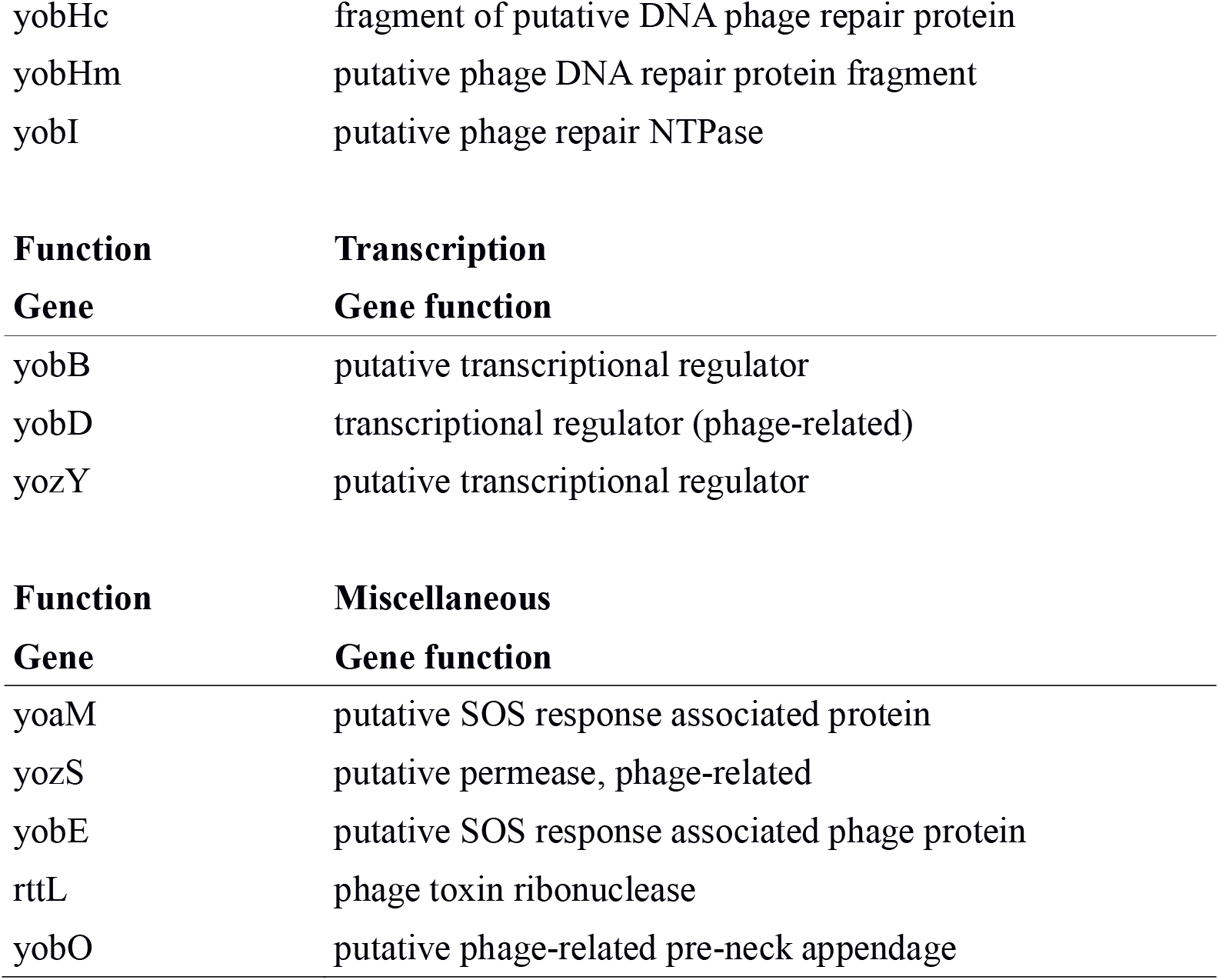
Categorization of function of annotated genes in prophage 6 of *Bacillus subtilis*.

### 3.3 Genetic repertoire of PBSX prophage in the genome of *Bacillus subtilis*

Table 5 shows the genetic repertoire of PBSX prophage in *Bacillus subtilis* 168. Specifically, the prophage genome comprises 41 genes, which is larger than the genome of prophage SPBeta and prophage 6 in *B. subtilis*. Classified as a defective prophage, PBSX prophage nevertheless still retain many prophage structural genes such as RNA polymerase, capsid portal protein, prophage terminase, and phage chromosome binding protein. Similar to other prophages in *B. subtilis* genome, prophage PBSX has many conserved hypothetical proteins whose function await elucidation.

**Table 5:**
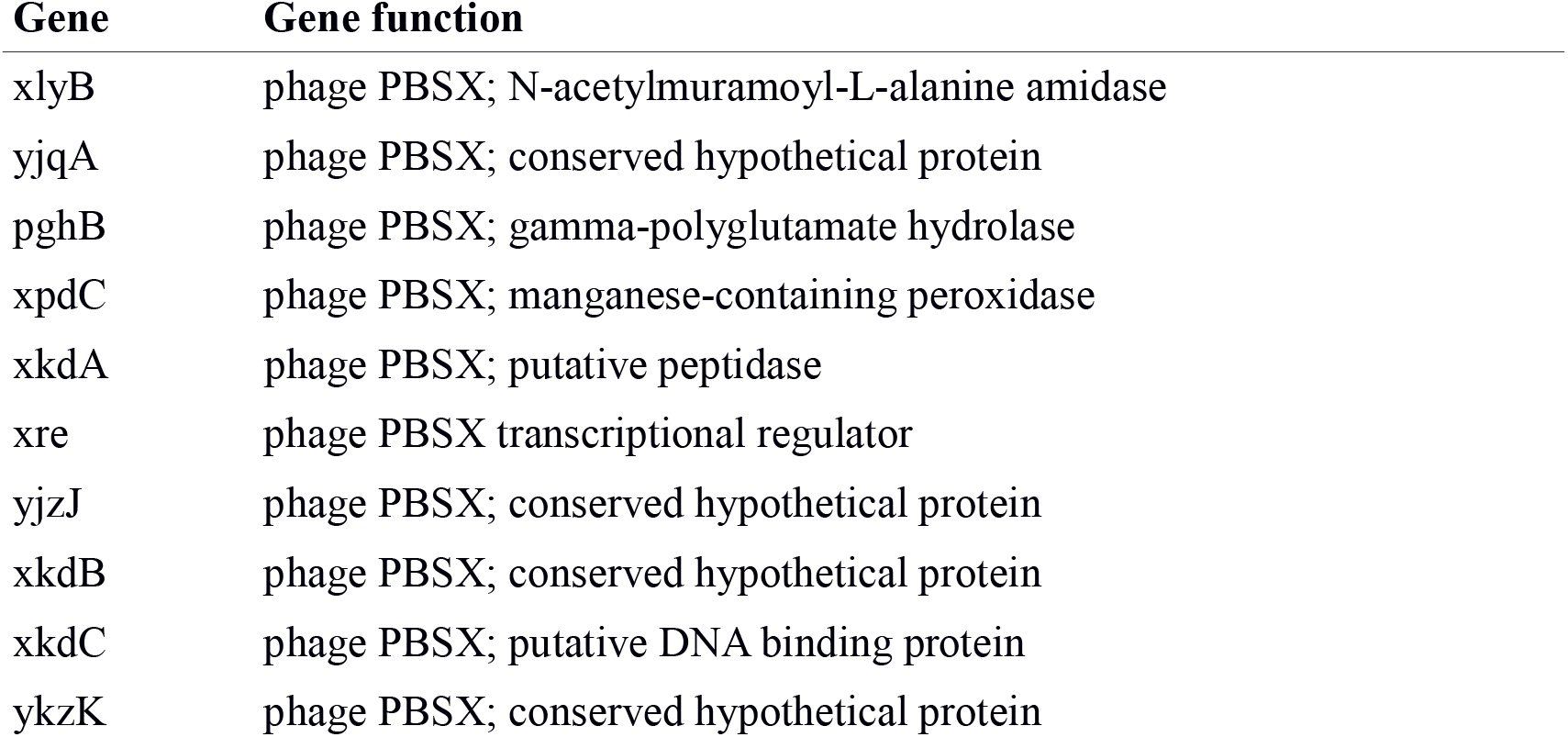
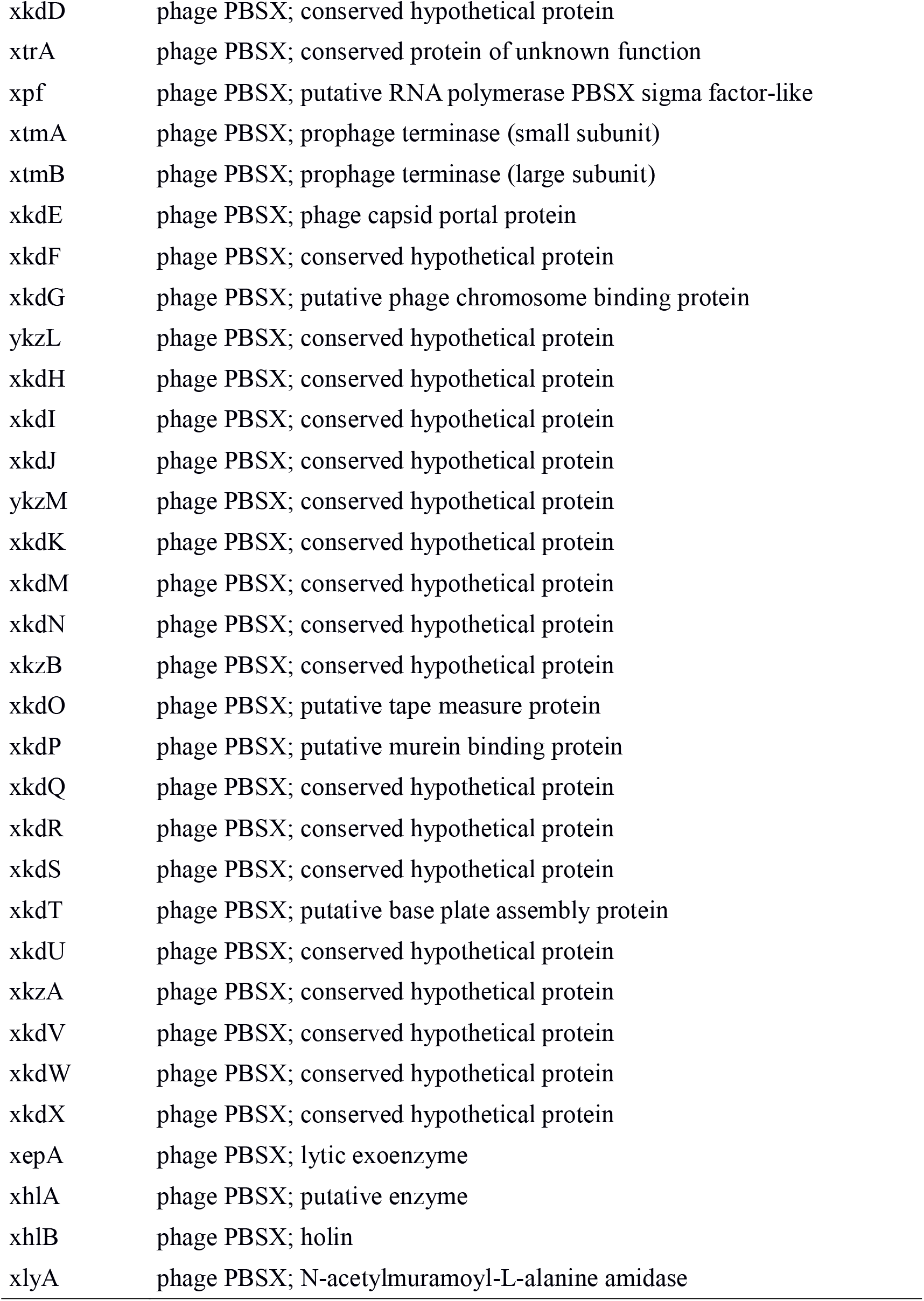
Genetic repertoire of PBSX prophage in *Bacillus subtilis* 168.

Categorization of genes in the PBSX prophage based on function reveals distinct functional categories in DNA replication, transcription, structural, and lytic phase (Table 6). Although PBSX is known to be a defective prophage in the *B. subtilis* virology community, it nevertheless still retains key genes in various important aspects of a prophage life cycle. For example, in the area of DNA replication, PBSX still retains DNA binding protein and phage chromosome binding protein. While these two genes may not be sufficient for orchestrating the entire DNA replication process, it indicates that host transposase and transposon activity are unable to completely destroy the prophage’s ability to replicate. Next, PBSX still retain at least two genes regulating transcription. These genes are critical as a prophage must possess the ability to regulate transcriptional activity at a fine balance in order not to put undue stress on the expression of host genes, while ensuring that sufficient phage genes are expressed and proteins produced. In PBSX, the two transcription related phage genes that are retained are transcriptional regulator and putative RNA polymerase PBSX sigma factor, but, it is important to note that this is likely to be an incomplete list of prophage genes involved in transcriptional regulation.

**Table 6:**
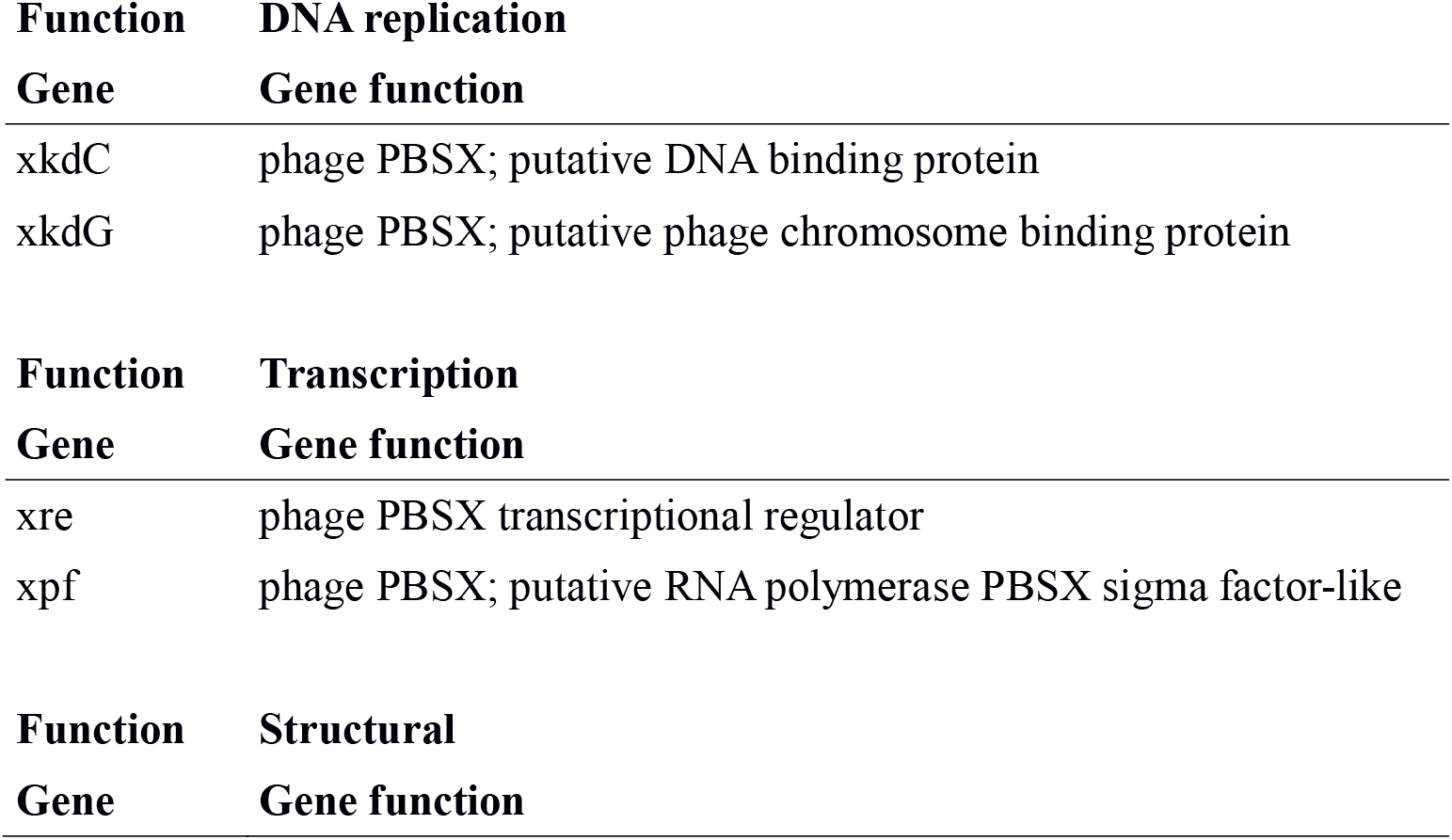
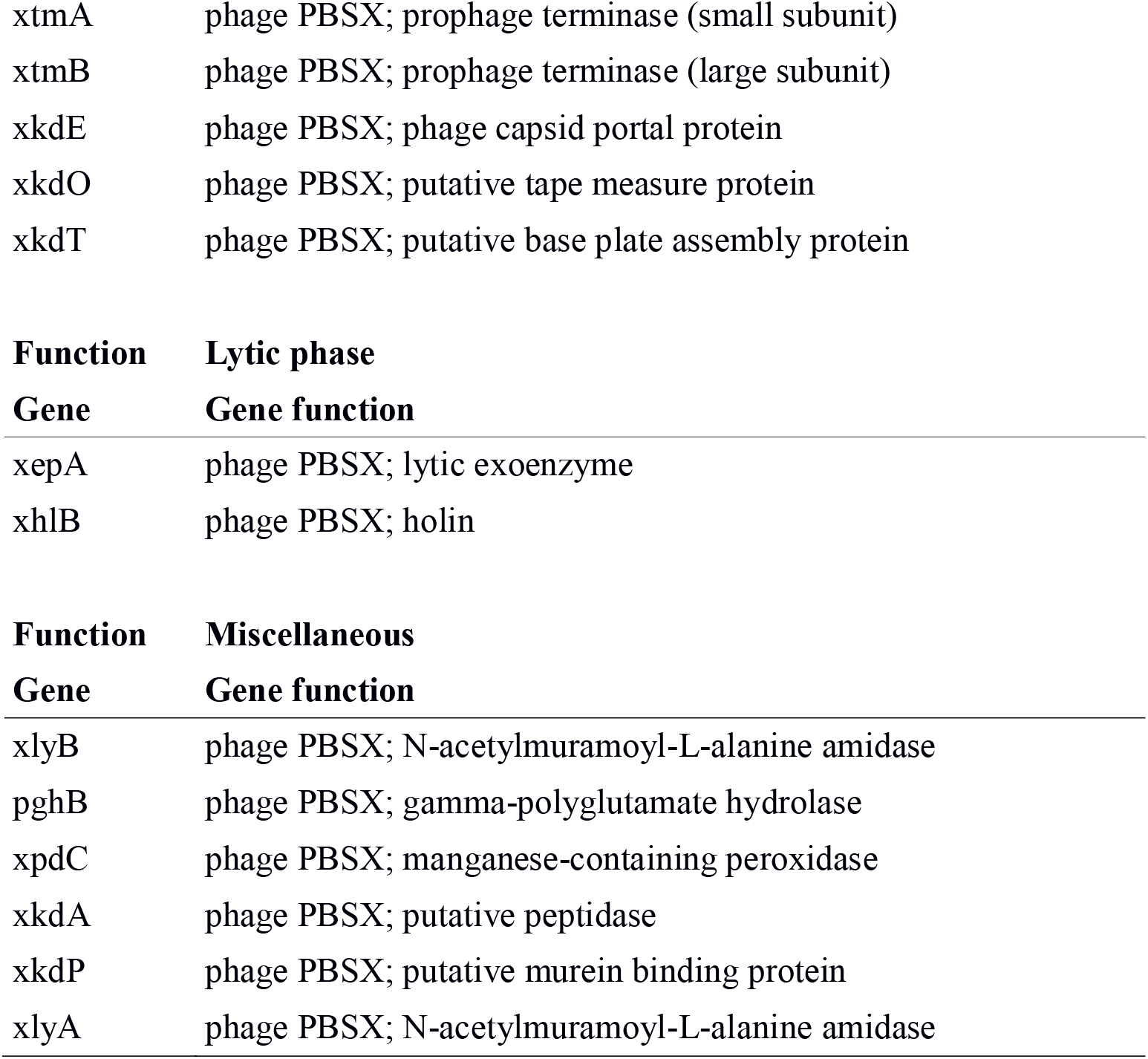
Functional categorization of genes in PBSX prophage.

| **Function** | **DNA replication** |
| --- | --- |
| **Gene** | **Gene function** |
| xkdC | phage PBSX; putative DNA binding protein |
| xkdG | phage PBSX; putative phage chromosome binding protein |
| **Function** | **Transcription** |
| **Gene** | **Gene function** |
| xre | phage PBSX transcriptional regulator |
| xpf | phage PBSX; putative RNA polymerase PBSX sigma factor-like |
| **Function** | **Structural** |
| **Gene** | **Gene function** |
| xtmA | phage PBSX; prophage terminase (small subunit) |
| xtmB | phage PBSX; prophage terminase (large subunit) |
| xkdE | phage PBSX; phage capsid portal protein |
| xkdO | phage PBSX; putative tape measure protein |
| xkdT | phage PBSX; putative base plate assembly protein |
| xepA | phage PBSX; lytic exoenzyme |
| xhlB | phage PBSX; holin |
| xlyB | phage PBSX; N-acetylmuramoyl-L-alanine amidase |
| pghB | phage PBSX; gamma-polyglutamate hydrolase |
| xpdC | phage PBSX; manganese-containing peroxidase |
| xkdA | phage PBSX; putative peptidase |
| xkdP | phage PBSX; putative murein binding protein |
| xlyA | phage PBSX; N-acetylmuramoyl-L-alanine amidase |

Structural genes are important for enabling prophages to self-assemble into virions. In this area, PBSX prophage still retain important genes encoding proteins such as prophage terminase (small and large subunit), phage capsid portal protein, putative tape measure protein, and phage base plate assembly protein. Similar to the case for DNA replication and transcription regulatory protein, PBSX is unlikely to retain the full complement of structural genes that defines the structure of the phage virion. For a prophage to enter the lytic phase of the phage life cycle, genes encoding proteins important to performing tasks during the lytic phase are important in addition to the key excisionase enzyme. For PBSX, it still retains lytic exoenzyme and holin as genes important to the lytic phase of the phage life cycle. Although incomplete, the genetic repertoire for lytic phase genes in PBSX prophage shows that transposase and transposon activity in the host genome is indeed random, and not specifically targeted at particular groups of genes. Finally, the genetic repertoire of PBSX prophage also contain a couple of host genes swapped in possibly by the action of transposase and transposons. These genes are categorized as miscellaneous in the functional analysis of the genetic complement of PBSX. A cursory analysis of the function of the swapped-in host genes suggests that they belong to different classes of enzymes and proteins, and therefore reaffirmed the random nature in which transposase and transposon swapped-in host genes for prophage genes.

### 3.4 Prophage 1 and Prophage 3 no longer qualify as prophages

Besides SPBeta, prophage 6 and PBSX prophage, the genome of *B. subtilis* 168 also contains two other genomic regions that suggest previous integration of prophage genes. These are prophage region 3 and prophage region 1. However, these prophage regions can no longer be characterized as prophages as too many of the prophage genes have been swapped out by host genes. Comparing the relatively intact nature of prophage genes in SPBeta, prophage 6 and PBSX prophage, the results suggest that the genomic region in which prophage 3 and prophage 1 integrated into may be favourable for integration of transposon or possess specific sequence recognition motifs for transposases.

### 3.5 Expression level of prophage genes in *B. subtilis*

Table 7 shows a tabulation of the most highly expressed prophage genes in *B. subtilis* 168. A quick glance of the data suggests an anomaly in the data that should first be dissected. Specifically, prophage genes should be transcriptionally silent in the *B. subtilis* genome. But, the RNA-seq transcriptome data of the organism shows that many of the prophage genes are transcriptionally active at high level. Another 215 prophage genes register a transcription count of 5 or less, and 82 prophage genes were not expressed. From a theoretical perspective, prophage genes with no expression level may be true phage genes controlled by phage promoters, and therefore could not be expressed in *B. subtilis*.

**Table 7:**
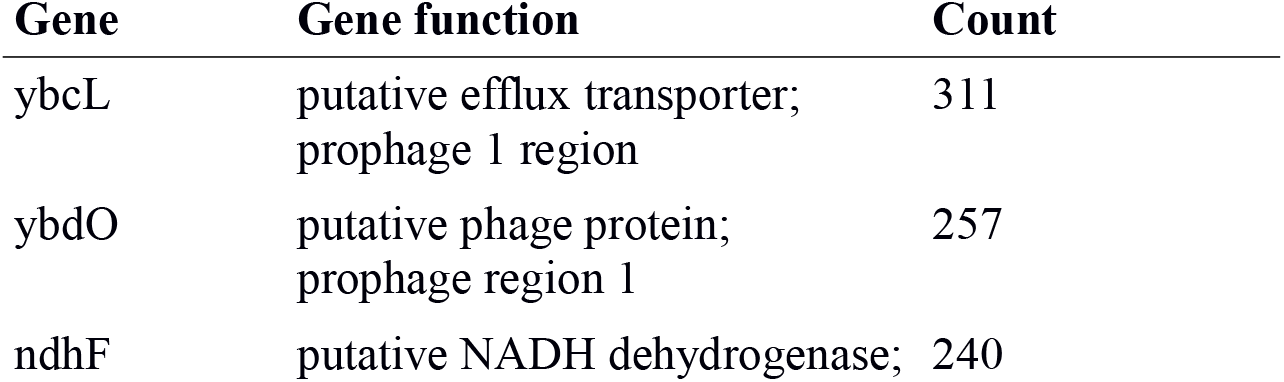
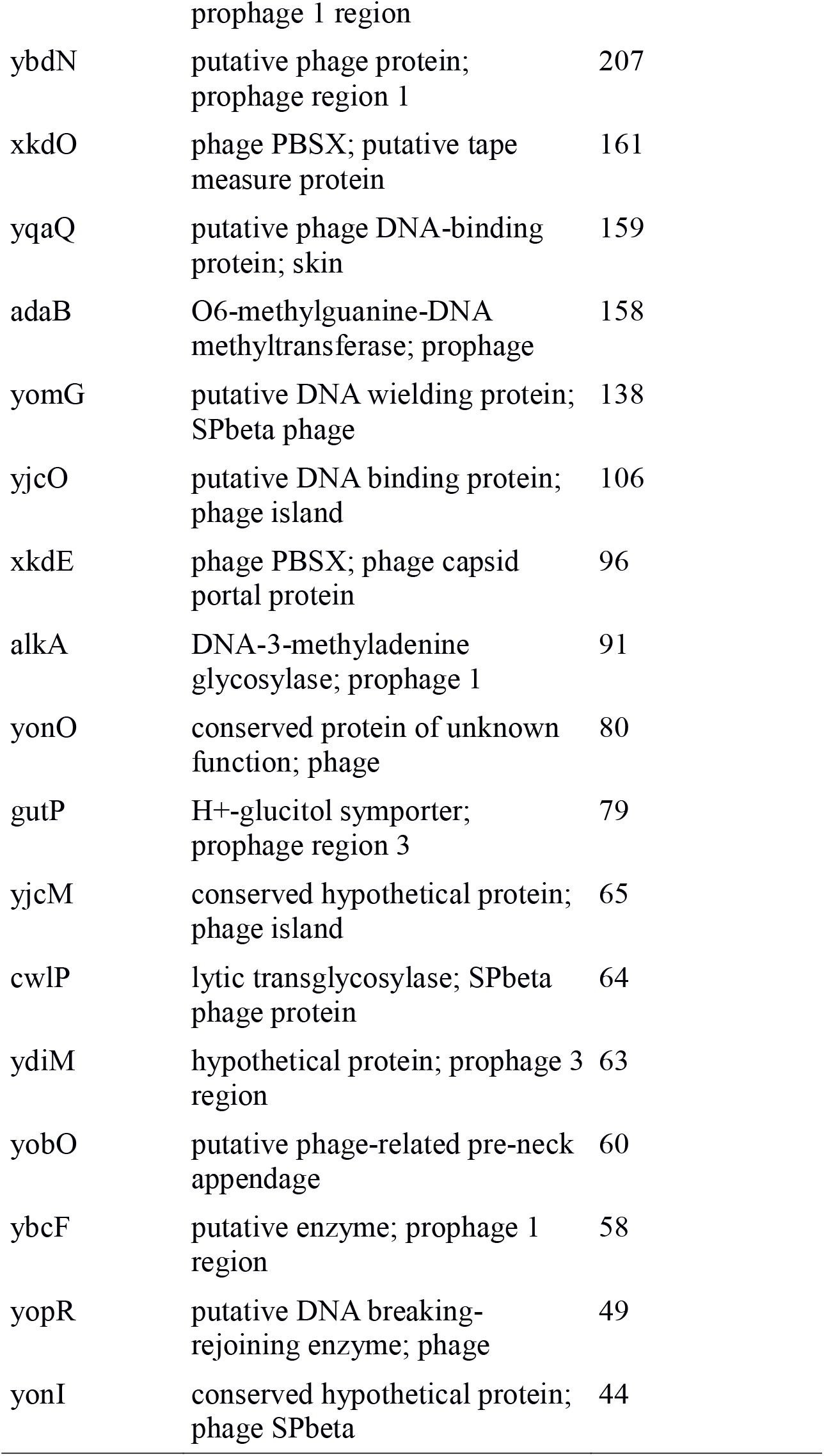
Expression level of highly expressed prophage genes in *B. subtilis* 168.

Presence of sizeable number of prophage genes with high expression activity suggests that these genes are controlled by promoters endogenous to *B. subtilis*. This could happen by way of transposase swapping in host promoters in place of prophage promoters in the prophage genome. Significant number of highly expressed prophage genes in *B. subtilis*, thus, further cements the role of transposase in recognizing unique genomic loci in prophage genome for splicing in of host genes and endogenous promoters. More importantly, many of the highly expressed prophage genes are putative phage genes, and not endogenous host genes swapped-in by transposase activity. The result highlights an evolutionary dance between prophage genome motif, and transposase activity endogenous to the host, whose provenance maybe possible close evolutionary relatedness and co-evolution between prophage and transposase and transposon outside of the context of the bacterial cell that serves as current host for the prophage and transposase. Such evolutionary relationships remain difficult to disentangle without the aid of further molecular evidence at the sequence level. But, one possible reason may be that transposase in *B. subtilis* might have “jumped out” of the prophage genome to reside at a separate location in the *B. subtilis* genome. Functional role of the transposase in the context of phages may be to aid the reshuffling of host genes into the phage genome to introduce mutations and augment the phage genetic repertoire for gaining a survival advantage against a multitude of restriction-modification, and CRISPR systems in bacterial system.

Analysis of the distribution of highly expressed prophage genes in different prophages further reveals that a significant percentage of the high expression genes are in prophage 1 region of *B. subtilis* (Table 8). Distribution of highly expressed prophage genes in other prophages are SPBeta prophage (30%), prophage 3 region (10%), and PBSX prophage (10%). Such a result confirms that prophage region 1 may harbour transposase recognition motifs that afford the swapping in of host genes or promoters, which, in the latter case, resulted in higher expression level of genes. But, this phenomenon is not restricted to prophage 1 region, and is also present in other prophage genome, but to a smaller extent.

**Table 8:** Distribution of highly expressed prophage genes in different prophages.

| Phage category | Number | Percentage abundance (%) |
| --- | --- | --- |
| Prophage 1 region | 6 | 30 |
| SPBeta prophage | 3 | 15 |
| Prophage 3 region | 2 | 10 |
| PBSX prophage | 2 | 10 |

Overall, RNA-seq transcriptome analysis reveals the paradoxical result that many prophage genes have high expression level. But, the result reveals that transposase in *B. subtilis* likely aid in swapping in of host promoters and genes into prophage genome, which helps expression of prophage genes. Evidence collected herein also suggests that transposase and transposons may be co-evolutionary mates with bacteriophages where transposase’s function is to aid the swapping of host genes into the phage genome to engender variation in the phage genetic repertoire that helps the phage gain a survival advantage in a harsh cellular environment policed by multiple bacterial restriction enzymes and CRISPR systems.

## 4. Conclusions

Throughout evolution, bacteriophage entered an evolutionary dance with their host bacterial species. Such co-evolution led to the development of lysogeny where bacteriophage integrated their genome into the host bacterial genome during times of environmental and nutritional stress. Analysis of the repertoire of prophage genes in the Gram-positive model bacterium, *B. subtilis* 168, reveals five distinct prophage regions: SPBeta, Prophage 6, PBSX, Prophage 3 region, and Prophage 1 region. The latter two prophages have been almost totally destroyed by transposase activity that swapped in host genes for phage genes. The remaining three distinct prophages are defective and likely extinct in the environment given that many of their key structural genes have been replaced by host genes likely due to transposase activity, and the prophage lost the ability to replicate its genome and exit the bacterial cell. RNA-seq transcriptome analysis further reveals the paradoxical result that many prophage genes enjoy high expression level. This points to the swapping in of host promoters into prophage genome to aid the expression of prophage genes as the only mechanism explaining the observations. Such a mechanism, in turn, highlights the evolutionary dance between host transposase and prophage, where transposase may be once a part of the prophage genome whose activity is to help swap in host genes to augment the genetic repertoire and diversity to help improve the survivability of the prophage against bacterial adaptive immune system and restriction-modification system.

## Conflicts of interest

The author declares no conflicts of interest.

## Funding

No funding was used in this work.

## Notes

### Competing Interest Statement

The authors have declared no competing interest.

